# Enteric viruses nucleic acids distribution along the digestive tract of rhesus macaques with idiopathic chronic diarrhea

**DOI:** 10.1101/2021.06.24.449827

**Authors:** Eric Delwart, David Merriam, Amir Ardeshir, Eda Altan, Yanpeng Li, Xutao Deng, J. Dennis Hartigan-O’Connor

## Abstract

Idiopathic chronic diarrhea (ICD) is a common clinical condition in captive rhesus macaques, claiming 33% of medical culls (i.e. deaths unrelated to research). Using viral metagenomics we characterized the eukaryotic virome in digestive tract tissues collected at necropsy from nine animals with ICD. We show the presence of multiple viruses in the *Parvoviridae* and *Picornaviridae* family. We then compared the distribution of viral reads in the stomach, duodenum, jejunum, ileum, and the proximal, transverse, and distal colons. Tissues and mucosal scraping from the same locations showed closely related results while different gut tissues from the same animal varied widely. Picornavirus reads were generally more abundant in the lower digestive tract, particularly in the descending (distal) colon. Parvoviruses were more abundant in the upper reach particularly in the stomach. In situ hydridization (ISH) of fixed tissues showed punctuated staining for both these RNA and DNA viruses in the distal colon. Parvovirus ISH staining was also detected in the stomach/duodenum/jejunum in distinct oval-shaped structures. Therefore, the location of enteric viral nucleic acid differed widely between different viral families and along the length of the digestive tract.

## Introduction

Idiopathic chronic diarrhea (ICD) in captive macaques is the most common cause of euthanasia that is not related to medical research in primate research centers, impacting the health of the animals and reducing the availability of healthy non-human primate (NHP) for biomedical research ^1-5^. Clinically ICD is seen as persistent or recurring non-bloody diarrhea with microscopic ulcers in the colon ^2,3,6-8^. The mucus layer is also thickened, with shortened crypts showing a very high level of lymphocyte and plasma-cell infiltration, typical signs of a response to viral infections. Affected animals were tested negative for enteropathogenic agents, including parasitic worms and bacteria (*Shigella, Campylobacter, Yersinia, Salmonella*, and C*lostridium difficile*) and are non-responsive to antibiotic treatments. These clinical observations point to viral infections as possible causes of ICD. This condition resembles human ulcerative colitis, a subset of human inflammatory bowel disease (IBD) affecting the colon and rectum (distinct from IBD’s other major form, Crohn’s disease, which affects the entire GI tract)^5,9^.

We recently described a number of viruses in feces of captive rhesus macaques with and without ICD^10^ as well as in healthy captive and wild cynomolgus macaques^11^. In captive rhesus macaques, numerous picornaviruses and parvoviruses were characterized as well as less prevalent adenovirus and sapelovirus^10^. Several enteroviruses were weakly associated with acute resolving diarrhea and ICD^10^.

Here we analyzed the distribution of viral reads along the digestive tracts of 9 rhesus macaques that were necropsied following diagnosis of ICD. Different gut tissues and scrapings of their surfaces were analyzed for viral nucleic acids following enrichment of virus-like particles associated nucleic acid, followed by non-specific amplification of DNA and RNA and next-generation sequencing. Fixed gut tissues were also hybridized with viral probes to identify their anatomical location of viruses along the digestive tract.

## Material and method

### Sample collection

The California National Primate Research Center (CNPRC) is an Association for Assessment and Accreditation of Laboratory Animal Care (AAALAC) accredited facility where animals are cared for in accordance with the USDA Animal Welfare Act and regulations and the *Guide for the Care and Use of Laboratory Animals*. Animal Welfare Assurance on file with the Office of Laboratory Animal Welfare (OLAW). The Assurance Number is D16-00272(A3433-01). Additionally, all procedures in this study were approved by the Institutional Animal Care and Use Committee (IACUC) of the University of California, Davis. All animals were born and housed at the CNPRC. Animals were provided with species-appropriate environmental enrichment, fed chow twice daily (LabDiet Monkey Diet 5047, Purina Laboratory, St Louis, MO), offered water without restriction through automatic watering devices, and supplemented with vegetables biweekly.

All rhesus macaques were submitted to necropsy due to the primary problem of idiopathic chronic diarrhea (ICD). Animals were considered to have ICD based on the following criteria: greater than 45 days of recorded diarrhea and/or greater than 3 hospitalizations for treatment of nonpathogenic diarrhea in the previous 6 months, tested negative for bacterial and parasitic pathogens in the previous 6 months, and no etiology for diarrhea identified during microscopic analysis of tissues or at the time of necropsy.

### Necropsy

All animals underwent complete necropsies, including cultures of small intestinal and large intestinal contents, and bile for *Campylobacter spp*., *Shigella flexneri, Yersinia* species, and *Escherichia spp*., as well as fecal examination by direct microscopy and IFA for *Giardia* and *Cryptosporidium*. The intestinal tract and any gross lesions identified at the time of necropsy were examined microscopically. Routine sections included stomach, duodenum, jejunum, ileum, proximal colon, transverse colon, and distal colon. Samples were stored at -80°C until use.

### Metagenomic analysis

Tissue and mucosal scraping samples from the stomach, duodenum, ileum, jejunum, proximal colon, transverse colon, and distal colon from nine animals were processed (a total of 126 samples) for metagenomic analysis. Tissues and mucosal scrapings were homogenized with a hand-held rotor in 1ml of PBS buffer with 100ul of zirconia beads and then rapidly frozen on dry ice and thawed five times before centrifugation at 9,000 rpm for 5 min. The supernatants were passed through a 0.45 µm filter (Millipore, Burlington, MA, USA) and filtrates digested for 1.5 hours at 37°C with a mixture of nuclease enzymes consisting of 14U of Turbo DNase (Ambion, Life Technologies, USA), 3U of Baseline-ZERO (Epicentre, USA), 30U of Benzonase (Novagen, Germany) and 30U of RNase One (Promega, USA) in 1x DNase buffer (Ambion, Life Technologies, USA). 150 ul was then extracted using the MagMAX^™^ Viral RNA Isolation kit (Applied Biosystems, Life Technologies, USA) and nucleic acids were resuspended in 50 µl water with 1 ul RiboLock RNAse inhibitor. 11 µl of nucleic acids were incubated for 2 min at 72^°^C with 100 pmol of primer A (5’GTTTCCCACTGGANNNNNNNN3’) followed by a reverse transcription step using 200 units of Superscript III (Invitrogen) at 50^°^C for 60 min with a subsequent Klenow DNA polymerase step using 5 units (New England Biolabs) at 37^°^C for 60 min. cDNA was then amplified by a PCR step with 35 cycles using AmpliTaq Gold™ DNA polymerase LD with primer A-short (5’GTTTCCCACTGGATA3’) at an annealing temperature of 59^°^C. The random amplified products were quantified by Quant-iT™ DNA HS Assay Kit (Invitrogen, USA) using Qubit fluorometer (Invitrogen, USA) and diluted to 1 ng of DNA for library input. The library was generated using the transposon-based Nextera™ XT Sample Preparation Kit using 15 cycles (Illumina, San Diego, CA, USA) and the concentration of DNA libraries was measured by Quant-iT™ DNA HS Assay Kit. The libraries were pooled at equal concentration and size selected for 300 bp – 1,000 bp using the Pippin Prep (Sage Science, Beverly, MA, USA). The library was quantified using KAPA library quantification kit for Illumina platform (Kapa Biosystems, USA) and a 10 pM concentration was loaded on the MiSeq sequencing platform for 2×250 cycles pair-end sequencing with dual barcoding.

### Bioinformatics

Human and bacterial reads were identified and removed by comparing the raw reads with human reference genome hg38 and bacterial genomes release 66 (collected from ftp://ftp.ncbi.nlm.nih.gov/blast/db/FASTA/, Oct. 20, 2017) using local search mode. Low quality sequence ends below Phred 30 scores, as well as adaptor and primer sequences were trimmed by using VecScreen^12^. The remaining reads were de-duplicated if base positions 5 to 55 were identical with one random copy retained. The reads were then de novo assembled by Ensemble Assembler^13^. Assembled contigs and all singlet reads were aligned to an in-house viral protein database (collected from ftp://ftp.ncbi.nih.gov/refseq/release/viral/, Oct. 20, 2017) using BLASTx (version 2.2.7) using E-value cutoff of 0.01. The significant hits to the virus were then aligned to an in-house non-virus-non-redundant (NVNR) universal proteome database using DIAMOND (10) to eliminate the false viral hits. Hits with more significant (lower) E-value to NVNR than to the viral database were removed. Remaining singlets and contigs were compared to all eukaryotic viral protein sequences in GenBank’s non-redundant database using BLASTx [23]. The genome coverage of the target viruses was further analyzed by Geneious R11.1.4 software (Biomatters, New Zealand).

### Enterovirus typing

Enterovirus contigs were typed using the genotyping tool https://www.rivm.nl/mpf/typingtool/enterovirus/.

### Quantitation of viral reads

Genetic variation was observed between homologous sequences belonging to the same enterovirus type. With the goal of assessing the viral abundance of each viral types, closely related contigs (of the same viral type) were concatenated (with 100 bp ‘N’ spacers to prevent reads aligning to the areas where the contigs are joined). These contigs of contigs were then used to find matching reads using the nucleotide aligner program Bowtie2 with the seed length parameter set at 30, and the alignment considered a hit if the read identity to the reference concatamer was greater than 95%. A total of 11 concatenated contigs representing the most commonly detected viruses were generated. For normalization (to account for the variable number of total reads generated from different samples) the number of matching read hits were converted to reads per million total reads (RPM). Comparing the distribution of RPM to different viruses in different tissues along the gut was done using one-way ANOVA (Kruskal-Wallis test).

### In situ hybridization using RNAScope

Tissues were fixed in 10% neutral buffered formalin between 3 days and 12 weeks prior to being routinely processed, embedded in paraffin, and sectioned at 5µm. One section of each animal tissue stained with RNAscope was also stained with Mayer’s hematoxylin and eosin-Y. RNA scope was done on a subset of animals, and probes were designed using the metagenomics data of those viruses of interest. Advanced Cell Diagnostics (ACD) (Newark, California, USA) generated four custom probes-two probes targeting enterovirus 19, one probe targeting erythroparvovirus, and one probe targeting a protoparvovirus. Paraffin-embedded tissues were stained using ACD’s RNAScope Fluorescent Multiplex Kit V2 following the manufacturer’s instructions. An equivalent steamer was used instead of the recommended brand in the protocol. TSA resolution was performed using TSA Cy5 from PerkinElmer, followed by 1 minute DAPI incubation and mounting with ProLong Gold. Slides were allowed to cure for at least 24 hours before image capture. Images were captured on an Evos2 Auto Fluorescent System. Samples were analyzed on FIJI (Fiji Is Just ImageJ) and utilizing FIJI’s built-in macro language for automated segmentation, deconvolution, and quantification.

## Results

The goal of this study was to characterize the eukaryotic viruses in different gut tissues of macaques suffering from ICD. Nuclease-resistant (viral particle-associated) nucleic acids were enriched from frozen/thawed tissue biopsies and from matching scraped mucosae acquired during necropsies. A total of seven necropsied tissues, plus mucosal scrapings from the same tissues, from each of nine animals with ICD were analyzed separately and the raw data deposited in GenBank (PRJNA607332). Illumina MiSeq runs were therefore used to analyze 126 samples, yielding a total of 78,985,006 reads, with a range of 19,724 to 2,571,490 reads and an average of 636,975 reads per samples. A total of 74 contigs greater than 500 bp were generated and grouped into 11 concatamers representing 11 viruses from three different viral families, *Picornavirudae, Caliciviridae*, and *Parvoviridae* (Table 1). Further information on the % sequence identity of each contigs are in supplemental data (Table S1). The concatenated sequences can be found in supplemental data (Dataset S1).

**Table 1.** The number of contigs per viral species included in the concatamers used for virus read quantitation using Bowtie.

| family | genus | species | type | No. of contigs |
| --- | --- | --- | --- | --- |
| *Caliciviridae* | Sappovirus | Sapporo virus |  | 2 |
| *Picornaviridae* | Enterovirus | A | SV19/EV-A122 | 23 |
| *Picornaviridae* | Enterovirus | A | SV46/EV-A124 | 5 |
| *Picornaviridae* | Enterovirus | A | EV-A92 | 9 |
| *Picornaviridae* | Enterovirus | J | EV-J103 | 9 |
| *Picornaviridae* | Sapelovirus | B |  | 7 |
| *Parvoviridae* | Bocaparvovirus | Unclassified |  | 1 |
| *Parvoviridae* | Protoparvovirus | Unclassified | WUHARV | 2 |
| *Parvoviridae* | Chaphamaparvovirus | Unclassified |  | 8 |
| *Parvoviridae* | Dependoparvovirus | Adeno-associated virus 11 |  | 2 |
| *Parvoviridae* | Erythroparvovirus | Unclassified |  | 4 |

These 11 contiguous viral “probes” were then used to quantify matching viral reads in each library (materials and methods). The results are shown as viral reads per million total reads (RPM) (Fig 1). Each animal showed a unique pattern of viruses in tissues and mucosal scrapings (Figure 1). The viral reads from tissues and mucosal scraping were generally although not always similar. For example in animal 45629 two viruses predominated which were mainly detected in the ileum tissue biopsy and corresponding mucosal scraping.

**Figure 1.**
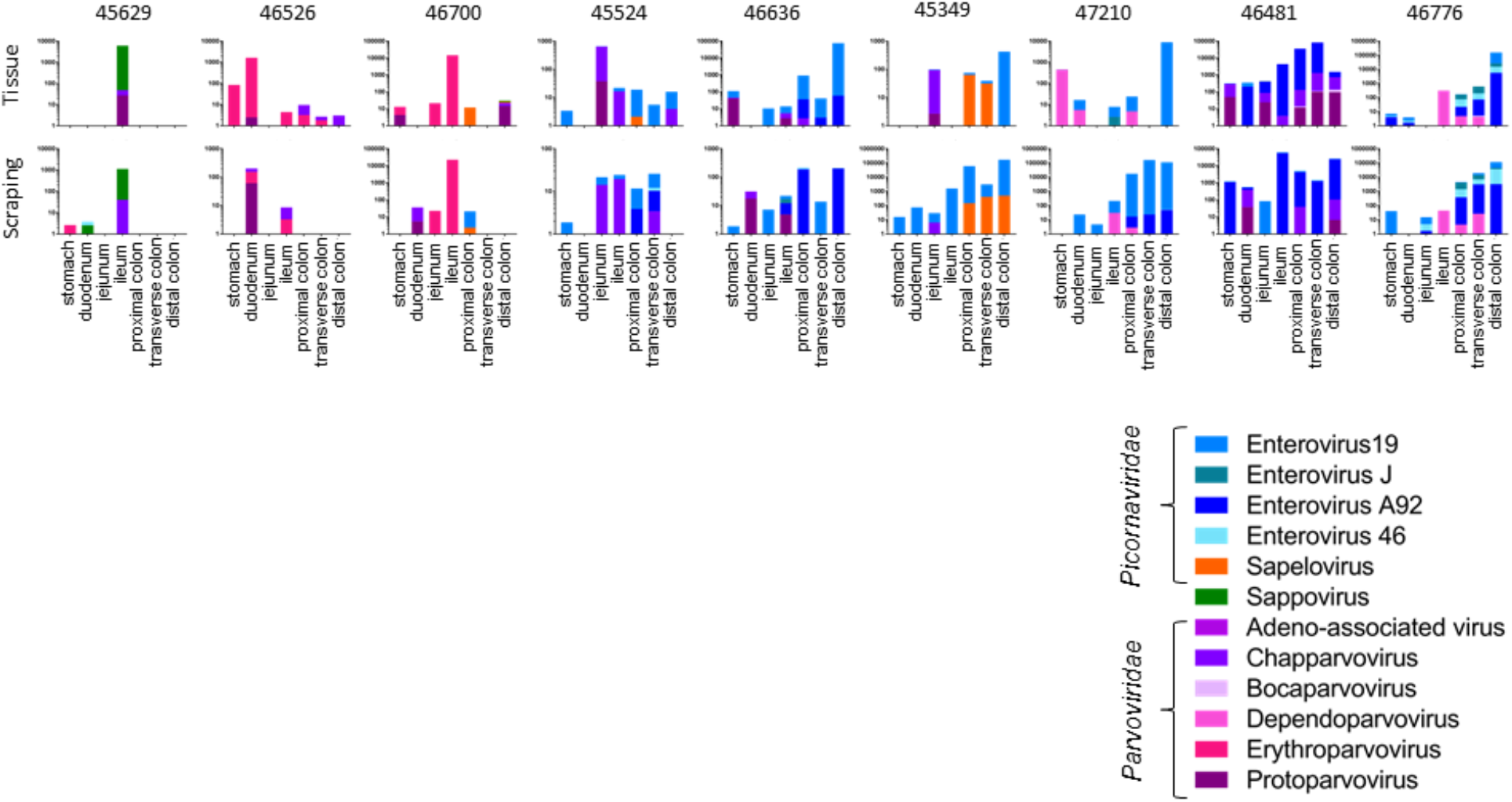
Viruses detected and their abundance in tissue and mucosal scrapings from the stomach, duodenum, jejunum, ileum, proximal colon, transverse colon, and distal colon of 9 macaques with ICD.

The following viruses were detected in the 9 ICD animals. In the *Picornaviridae* family (enterovirus B species) we detected enterovirus 19 and 46 in seven animals each and enterovirus A92 in five macaques. Enterovirus J was found in two and sapelovirus in three animals. Sapovirus in the *Caliciviridae* family was found in one animal. In the *Parvoviridae* family we found chapparvovirus in seven, protoparvovirus in six, erythroparvovirus in three, dependoparvovirus in two, and bocaparvovirus in one animal. The viral pattern observed by viral metagenomics was generally concordant between the biopsied tissues and the overlying mucosal scrapings indicating that the viruses within the mucosal layer largely reflected those in the underlying tissues. The viral abundance of the different viruses from these animals were then plotted by anatomical location. A higher number of reads was observed in the lower intestines which were dominated by enteroviruses (Figure 2).

**Figure 2.**
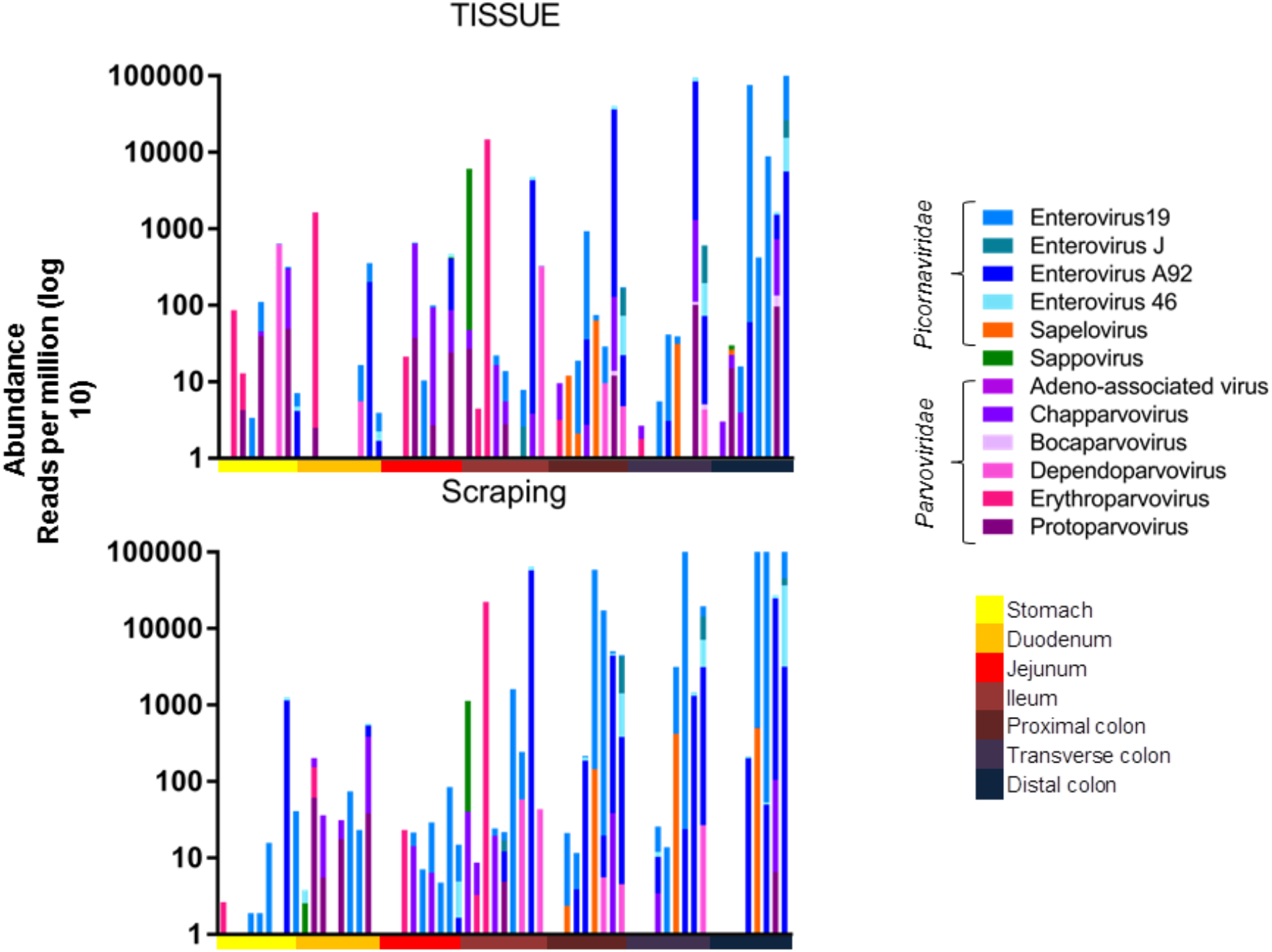
Abundance of viral reads along length of the digestive tract. Top panel: Abundance of eukaryotic viral reads from tissues along the intestinal tract for all animals. Bottom panel: Abundance of eukaryotic viral reads from mucosal scrappings along the intestinal tract for all animals. Blue shading represents different enteroviruses, while shades of pink or purple represent different *parvoviruses*.

A one-way ANOVA (Kruskall-Wallis test) was used to compare the total viral read abundance of the tissues and mucosal scrapings along the gastrointestinal tract, resulting in a p-values of p<0.0001 and p=0.0017, respectively. The viral RPM values therefore increased towards the end of the digestive tract for both tissues and mucosal scrapings. The read abundance was also plotted after amalgating reads of the different picornaviruses and different parvoviruses (Fig 3).

**Figure 3.**
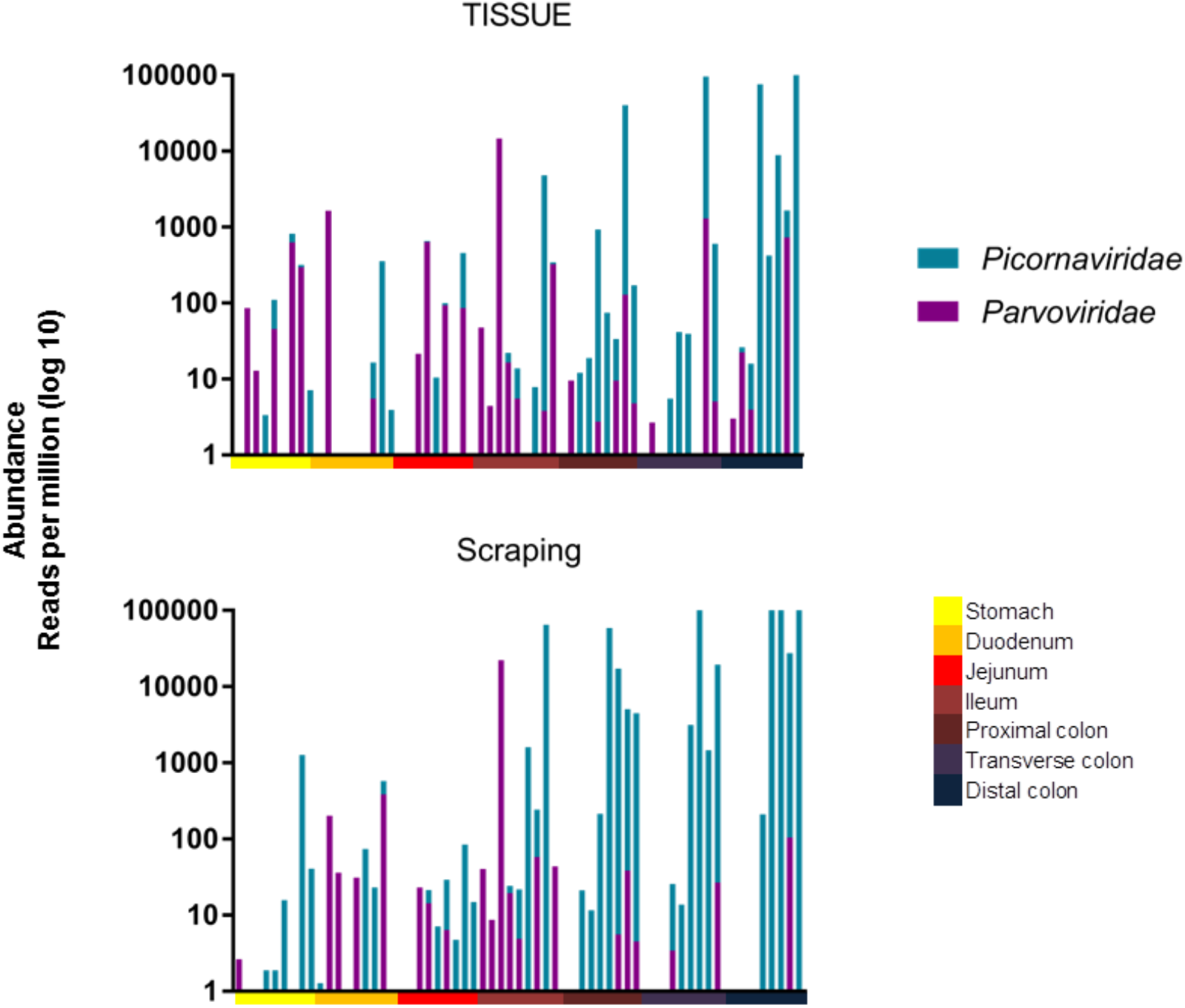
Abundance of picornavirus and parvovirus reads along the length of the digestive tract. Top panel: Abundance of eukaryotic viral reads from tissues along the intestinal tract for all animals. Bottom panel: Abundance of eukaryotic viral reads from mucosal scraping along the intestinal tract for all animals. Shades of blue represent different enteroviruses, while shades of pink or purple represent different *parvoviruses*.

The same statistical test was used to show that there was a significant difference in their distribution with parvoviruses being found further up the digestive tract than picornaviruses when analyzing mucosal scraping (P=0.0003), but only a trend when analyzing tissue biopsies (p=0.0856).

To assess the similarity, measured by viral presence and abundance, of viruses at each location of the gastrointestinal tract, a Bray-Curtis dissimilarity matrix was created and plotted as a multidimensional scale (MDS) (Fig 4). The MDS plot shows that the viral reads in the large intestine (blueish colors) cluster differently than those in the stomach and small intestine (reddish colors). Based on the abundance of different viruses, this clustering was likely driven by the higher abundance of picornaviruses in the lower intestinal tract.

**Figure 4.**
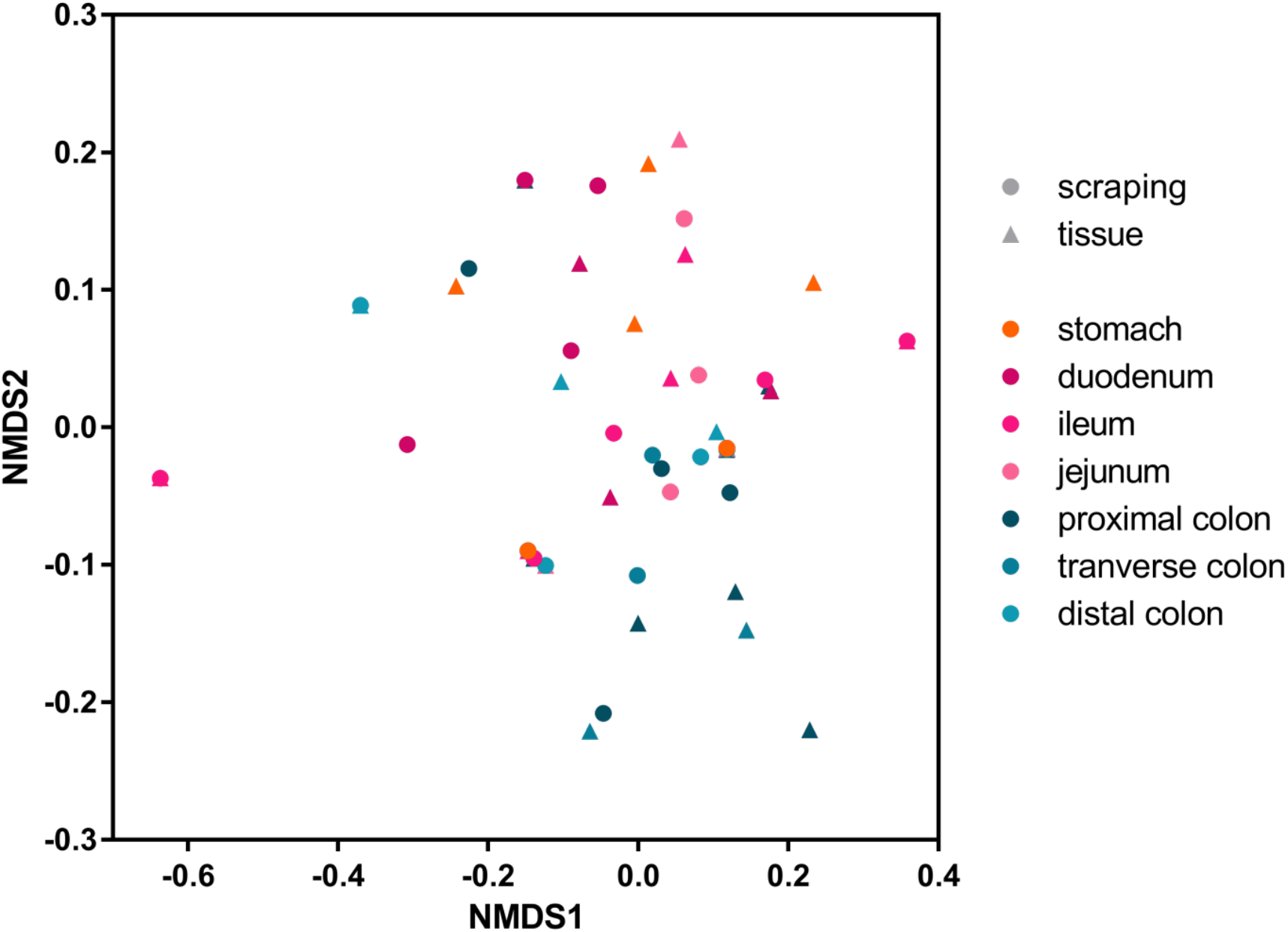
MDS plot of a Bray-Curtis dissimilarity matrix. The large intestine tissues are represented in shades of blue, the small intestine tissues are in shades of pink, and the stomach is in orange. Circles represent mucosal scrapings, while triangles represent tissue samples.

In situ hybridization was next used to locate anatomical sites of viral RNA by annealing fluorescently labeled RNAScope probes to fixed gut tissues. Parvovirus RNA was used to probe different tissues from the digestive tract of two animals and enterovirus RNA to probe the same tissues from three animals. Images of positive microscopic fields in different tissues are shown (Fig 5A, C).

**Fig 5.**
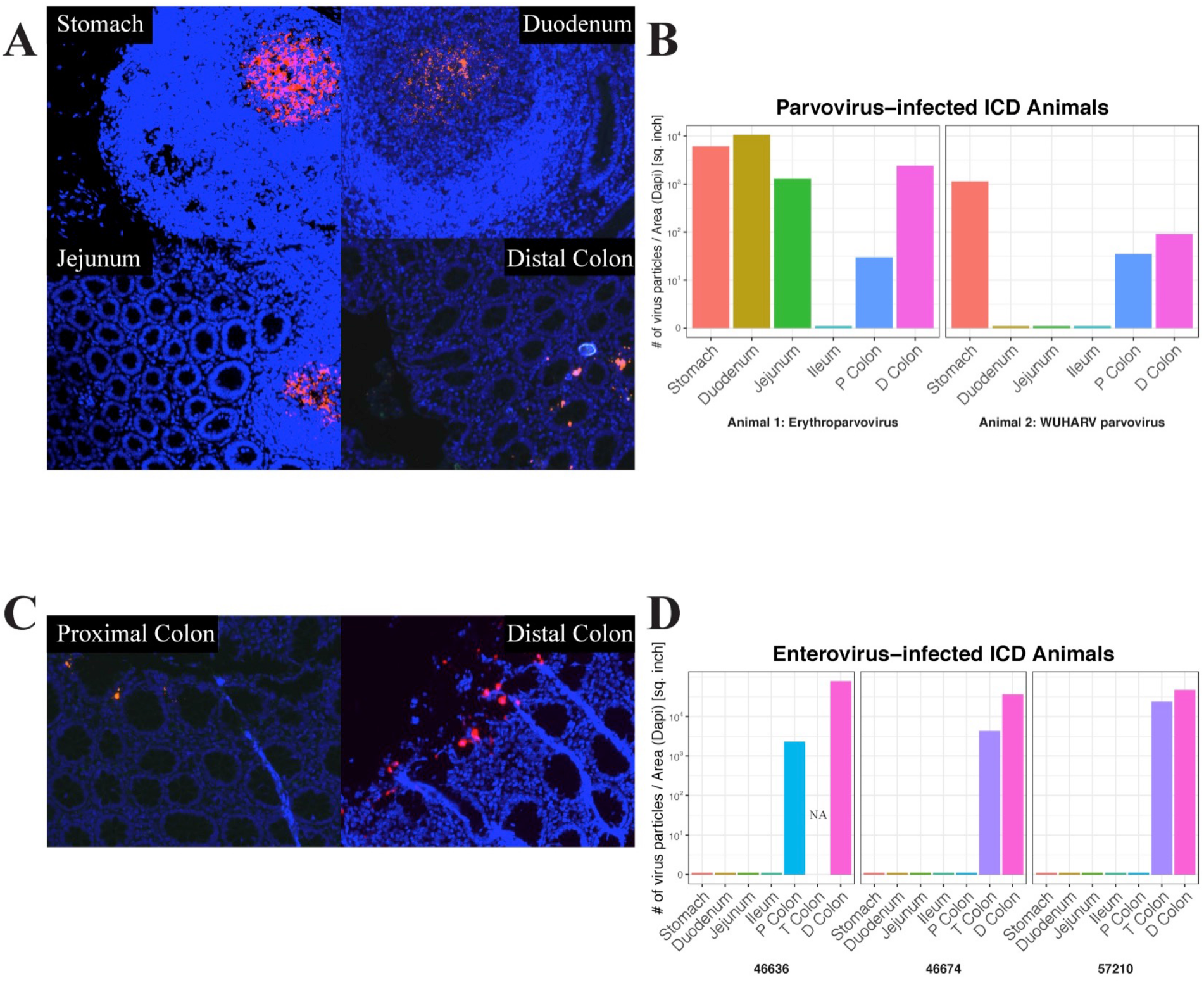
Representative positive microscope fields of different gut tissues hybridized with RNAScope viral probes. A. Erythroparvovirus probes hybridized with 46526 tissues. B. Quantitation of fluorescent signal of erythroparvovirus RNA in 46526 (left frame) and protoparvovirus RNA probe in 46700 (right frame). C. Enterovirus 19 probe hybridized with 46636 tissues. D. Quantitation of RNAScope signal for enterovirus 19 RNA in 46636 (left frame), 46774 (middle frame), and 57210 (right frame).

The signal distribution of erythroparvovirus RNA was restricted to oval-shaped structures in the stomach, duodenum, and jejunum which were reminiscent of Peyer’s patches. Distinct, more punctuated, staining was also seen in the distal colon (Fig 5A). When the erythroparvovirus ISH signal was quantified, it similarly showed high staining in the stomach, duodenum, and jejunum tissue and then in the lower transverse and distal colons (Fig 5B left frame). The distribution of protoparvovirus RNA in another animal showed a similar pattern of staining in the stomach and the lower colon tissues only (Fig 5B right frame). The distribution of enterovirus 19 RNA was also probed in three animals: 46636, 46776, and 57210 (Fig 5D). For enteroviruses only a punctuated distribution of fluorescent signal was detected in the proximal, transverse, and distal colons, i.e., exclusively in the lower digestive tract (Fig 4C).

Fluorescent signal quantitation therefore supported the visualized signal distribution as well as the results of the deep sequencing indicating greater viral signal in the upper digestive tract, particularly the stomach, for parvoviruses and in the lower tract, particularly the distal colon, for the picornaviruses.

## Discussion

We show here using viral metagenomics and in situ hybridization that the distribution of viral nucleic acids differs along the length of the digestive tract. Enterovirus reads were found at a greater frequency in lower parts of the colon than in the upper digestive tract. The converse was observed for parvovirus reads which were more common in the upper digestive tract. The viral reads from the mucosal surface scrapings largely tracked those of the underlying tissues. A subset of animals and viruses were also analyzed using in situ hybridization to locate viral nucleic acids in fixed tissues. The metagenomics and the ISH results were broadly concordant with strong erythroparvoviruse staining in the stomach, jejunum, and duodenum, and for protoparvovirus, in the stomach of another animal in structures evocative of Peyer’s patches. Parvovirus staining was also seen in the distal colon but in a more punctuated form, possibly reflecting viral particle production at the tip of villi structures.

A previous study of rhesus macaques with ICD reported that the lumen contents of the proximal colon, distal colon, and rectum showed more picornavirus reads than in the more upstream terminal ileum ^14^. Here picornavirus reads were more abundant in the colon, especially the terminal and distal colon while very few reads or RNA staining were detected in the ileum or further upstream in the digestive tract consistent with this prior study. This study also reported no difference in parvovirus reads distribution from the ileum down ^14^. Here we found parvovirus reads (and ISH staining) starting higher up in the digestive tract, including stomach, duodenum, and jejunum, anatomical sites not analyzed in the prior study. Parvovirus reads were also detected in the proximal and distal colon but not in the ileum. The distribution of parvovirus ISH signal in different locations was also highly distinct, with the terminal colon showing punctuated staining while the staining in the stomach/duodenum/jejunum was distributed in large oval structures. The more punctuated ISH signals seen in the lower intestine may reflect virus replication in highly permissive cells. The nature of these oval structure remains uncertain but could consist of gut-associated lymphoid tissues (GALT) in Peyer’s patches, where parvovirus particles may be presented to B lymphocytes.

We, therefore, show that in rhesus macaques with idiopathic chronic diarrhea, viral nucleic acids are unevenly distributed along the length of the digestive tract. The viral reads in mucosal scrapings largely reflect those in the underlying tissues. Parvovirus RNA was found in oval-shaped structures from the stomach to the jejunum, while both picornaviruses and parvoviruses RNA were also found in the lower part of the large intestine in a different, more punctuated, staining pattern.

## Acknowledgments

We thank E. Fahsbender for data generation and R. Bruhn for assistance for statistical analyses. This research was supported by NIH R01AI123376 grant for support.

